# Gene Therapy for Efficient Suppression of T-Type Channels in Treating Diabetic Neuropathy

**DOI:** 10.64898/2025.12.08.692919

**Authors:** D. Duzhyy, A. Dovgan, N. Hrubiian, N. Kononenko, A. Romanenko, N. Senchenko, E. Lukach, N. Voitenko, O. Hubar, P. Belan

## Abstract

Painful diabetic neuropathy (PDN), a chronic and often incurable syndrome, is one of the most common and unpleasant complications of diabetes. Effective clinical interventions for PDN are very limited and already developed approaches are characterized by lack of molecular or cellular target specificity and a short duration of therapeutic effects. Numerous investigations causally link upregulation of the Cav3.2 T-type Ca^2+^ channels in peripheral nociceptive neurons to painful symptoms of PDN. Here we suggest an approach to alleviate these symptoms based on implementation of virus-mediated cell-specific delivery of vectors expressing small hairpin RNAs (shRNAs). Processed by Dicer into specific small interfering RNA (siRNA), they would suppress the expression of T-type Ca^2+^ channels. In order to experimentally validate this approach, we have initially confirmed the ability of designed Dicer-substrate small interfering RNAs (DsiRNAs) to suppress expression of T-type channels in neurons of primary hippocampal and dorsal root ganglia (DRG) cultures. Target sequences of the effectively interfering DsiRNA were then used to design shRNAs and the coding sequences of shRNAs were cloned into the vector pAAV under U6 promotor. This plasmid was also proved to be effective in interference with expression of the T-type channels in the rat cultured DRG neurons. The expression cassette of this plasmid will be packed into AAV6 particles with tropism to unmyelinated fibers to suppress T-type channel expression in nociceptive DRG neurons and to alleviate painful symptoms of PDN.

## Introduction

According to the International Diabetes Federation, by 2035, one in ten people worldwide will have diabetes, with the majority living in low- and middle-income countries. Diabetic neuropathy is one of the earliest, most common, and most unpleasant complications of diabetes, occurring in approximately 60% to 70% of people with diabetes (Gooch and Podwall, 2004), of whom approximately 1/3 suffer from severe burning, electric shock-like, or stabbing pain. However, the awareness, understanding and treatment of diabetic neuropathy are limited, as its molecular mechanisms are still poorly understood, despite the increasing number of studies worldwide. Targeted basic research into the mechanisms of diabetic neuropathy and the development of new approaches based on them are urgently needed to accelerate the implementation of new treatment strategies for painful diabetic neuropathy (PDN).

PDN is associated with changes in the excitability of primary neurons of the dorsal root ganglia (DRG), the so-called nociceptors, and the efficiency of synaptic transmission between them and neurons of the dorsal horn of the spinal cord (Todorovic, 2016)(Bourinet et al., 2016)(Weiss and Zamponi, 2019). Low-threshold voltage-gated T-type calcium channels (T-channels), which are highly expressed in primary nociceptors (Rose et al., 2013), regulate both neuronal excitability (Duzhyy et al., 2015) and presynaptic glutamate release at their terminals (Jacus et al., 2012). Numerous pharmacological studies support the role of T-channels in primary nociceptors in the processing of pain signals. Thus, T-channels represent an attractive pharmacological target for the treatment of pain resistant to currently available analgesics (Todorovic, 2016)(Cai et al., 2021). The T-channel family consists of 3 isoforms: Cav3.1, Cav3.2, and Cav3.3. The Cav3.2 isoform is more highly expressed in nociceptive neurons than others and is currently the most promising target for pain treatment (Cai et al., 2021). Increased functional expression of Cav3.2 T-channels, leading to a significant increase in T-type calcium current (T-current), has been found in primary nociceptors isolated from animals with PDN (Latham et al., 2009)(Khomula et al., 2013)(Khomula et al., 2014). Moreover, selective local knockdown (Messinger et al., 2009) or pharmacological blockade of the Cav3.2 T-channels in vivo (Latham et al., 2009)(Obradovic et al., 2014) effectively attenuated thermal hyperalgesia in diabetic neuropathy in animals with type 1 and type 2 diabetes. Thus, it is most likely that in PDN of both types of diabetes, activation of different metabolic signaling cascades ultimately leads to modulation of the Cav3.2 T-channels in primary nociceptors (Joksimovic et al., 2020)(Ficelova et al., 2020). Regardless of the precise mechanisms of this modulation, increased Cav3.2 T-channel activity and increased corresponding T-currents in nociceptive DRG neurons are causally linked to the maintenance of PDN, making Cav3.2 T-channels a promising target for its treatment. Despite great efforts, the development of highly selective Cav3.2 inhibition still remains a major challenge, in particular due to the structural similarity of T-channel isoforms. Another problem preventing the widespread use of Cav3.2 inhibition for the treatment of pain is the broad expression of Cav3.2 in various tissues; thus, systemic inhibition of Cav3.2 would lead to significant side effects. To date, there are several organic T-channel inhibitors and antisense oligomeric deoxynucleotide constructs that have shown analgesic effects in diabetic neuropathy when injected intrathecally or systemically (Todorovic, 2016)(Cai et al., 2021). Many other novel approaches to indirectly target Cav3.2 channels are currently under development (Cai et al., 2021). However, all of the approaches that have been developed are characterized by a lack of specificity in targeting at the molecular or cellular level, complex or significantly invasive procedures, insufficient degree of inhibition, and short duration of therapeutic effects. Thus, they are not ideal for the treatment of PDN, whereas targeted gene therapy represents a novel and promising therapeutic strategy, especially when combined with simple and minimally invasive delivery methods for blocking agents. In this study we used knowledge of the rat Cav3.2 cDNA sequence to design DsiRNA and shRNA-coding expression cassettes cloned into the pAAV vector under U6 promotor, that effectively blocked functional expression of the Cav3.2 T-type channels in primary cultured hippocampal and DRG neurons. shRNA-coding expression cassettes is ready to be packed into AAV6 particles for specific delivery of these cassettes into nociceptive DRG neurons *in vivo* and for alleviating the symptoms of PDN.

## Materials and Methods

All animal care and handling were done in accordance with protocols of the Animal Care and Use Committee at Bogomoletz Institute of Physiology and conformed to the NIH Guide for the Care and Use of Laboratory Animals and the Public Health Policy. An approval protocol of Bioethics Committee of Bogomoletz Institute of Physiology No 3/19 from 04/02/2019.

### Hippocampal primary cell cultures

Neurons were obtained from newborn Wistar rats (age postnatal day 0–1; 26 animals of both sexes for the whole work) killed via rapid decapitation without anaesthesia. All rats were from the vivarium of Bogomoletz Institute of Physiology. Hippocampi of the rats were enzymatically dissociated with trypsin. The cell suspension (initial density of 3–5×10^5^ cells per cm3) was plated on glass coverslips coated with laminin and poly-L-ornithine (Thermo Fisher Scientific, USA). Cells were maintained in feeding solution consisting of minimal essential medium, 1% horse serum 1% N2 supplement and 2% of B27 supplement (Thermo Fisher Scientific, USA) in a humidified atmosphere containing 5% CO2 at 37 °C as previously described (Dovgan et al., 2010).

### Primary cultures of DRG neurons from L4-L6 DRGs

Prior to ganglia isolation, glass coverslips in 12-well culture plates (well diameter ∼21 mm) were coated with 200 μl of an aqueous solution containing poly-L-lysine (0.4 mg/ml) and laminin (5 μg/ml). Coating was performed in a laminar flow hood, after which the plates were closed and placed in a CO_2_ incubator until use. This treatment facilitated neuronal attachment. In parallel, 1 ml of F12+ medium was conditioned in a CO_2_ incubator for 15 minutes, transferred to a sterile Eppendorf tube in the laminar flow hood, sealed, and placed near the dissection microscope to be used during ganglia isolation.

For isolation, rats weighing ∼100 g were anesthetized with an intraperitoneal injection of ketamine (200 μl) and xylazine (60 μl) diluted in 0.6 ml saline. Following decapitation, the dorsal skin was incised, the exposed spinal region was disinfected with 70% ethanol, and ribs with connective tissue were cut close to the spine. The vertebral wall was then cut horizontally on both sides, allowing the dorsal portion of the spine to be lifted while leaving the spinal cord attached to the ventral half. Under a dissecting microscope, the spinal cord was removed with forceps from the lateral recesses of the vertebral canal. DRG ganglia were identified as spherical structures within these recesses and were carefully dissected by pulling adjacent tissue with forceps, severing connective tissue and nerve roots, and transferring the freed ganglia into an Eppendorf tube containing F12+ medium supplemented with collagenase (5 mg/ml; Worthington, Type IV, cat. #LS004188). Tubes were disinfected externally with 70% ethanol and transferred to the laminar flow hood.

In the hood, ganglia were gently pipetted several times through a wide-bore tip to facilitate enzyme penetration, then transferred to a 35-mm Petri dish and incubated in a CO_2_ incubator for 2 hours. After incubation, ganglia were returned to an Eppendorf tube, and the collagenase solution was removed while leaving the ganglia at the bottom. They were washed three times with 1 ml of calcium- and magnesium-free HBSS (Hanks’ Balanced Salt Solution) that had been conditioned in a CO_2_ incubator for 15 minutes.

Following washes, ganglia were treated with 1 ml of trypsin solution (2.5 mg/ml; Worthington, cat. #LS003703) prepared in conditioned HBSS. The trypsin solution was mixed by gentle pipetting and transferred onto the ganglia, which were then pipetted several times through a wide-bore tip and incubated in a new 35-mm Petri dish for 30 minutes in a CO_2_ incubator. After trypsinization, ganglia were returned to an Eppendorf tube and washed twice with 1 ml of F12+ medium, followed by a third wash in which 1 ml of F12+ was added and retained.

To dissociate cells, ganglia were triturated by pipetting through progressively narrower sterile tips: three tips with cut ends to allow ganglia to pass freely, followed by a final uncut tip to achieve finer dissociation. The resulting cell suspension was plated at 150 μl per well onto 18-mm glass coverslips in 12-well plates. Cultures were maintained in F12+ medium in a CO_2_ incubator for approximately 24 hours prior to transfection.

### Transfection of hippocampal primary cultures

Hippocampal neurons were transfected after 13–17 days in culture using Lipofectamine 2000 transfection reagent essentially as described by the supplier (Invitrogen). All cultures were used for the experiments 2–3 days after transfection. Transfection success was 0.2–1.0%.

### Transfection of primary culture of DRG neurons

Primary cultures of dorsal root ganglion (DRG) neurons were prepared on glass coverslips placed in 12-well plates (well diameter ∼21 mm). Cells were seeded the previous day in F12 medium supplemented with fetal bovine serum (FBS) and gentamicin. On the day of transfection, the culture medium was removed and replaced with 0.5 ml of conditioned Neurobasal A medium containing gentamicin. Conditioning was performed by transferring the required volume of Neurobasal A medium into a 35-mm Petri dish and incubating it in a CO_2_ incubator for at least 15 minutes prior to use.

For transfection, 100 μl of transfection mixture was carefully dispensed onto each coverslip using a 200 μl pipette, after which the plates were incubated in a CO_2_ incubator for 4 hours. Following incubation, plates were transferred to a laminar flow hood, and the transfection mixture was removed from each well using a 1 ml pipette tip. Wells were replenished with 1 ml of conditioned F12 medium supplemented with FBS and gentamicin, using a fresh pipette tip for each well. Cultures were maintained in the CO_2_ incubator for 3 days before coverslips were removed and mounted in the recording chamber for electrophysiological experiments under an inverted microscope.

The transfection mixture for each well was prepared by combining 100 μl of jetOPTIMUS buffer with 0.6 μg of plasmid DNA in a 1.5 ml Eppendorf tube, followed by vigorous mixing on a vortex and brief centrifugation to collect the solution at the bottom. A 0.6 μl drop of jetOPTIMUS reagent was then placed on the wall of the tube above the buffer–DNA mixture, gently tapped so that the drop fell into the solution, and mixed several times on the vortex at maximum speed. The tube was again briefly centrifuged to collect the mixture, which was then allowed to stand at room temperature (23–25 °C) for 10 minutes before application to the neuronal cultures. All steps were performed at room temperature, and when mixtures were required for multiple wells, reagent volumes were scaled proportionally to the number of wells to be transfected.

### Electrophysiological recordings

Hippocampal neurons in primary cultures were visualized using inverted microscopes (IX70 or IX71; Olympus, Tokyo, Japan). Whole-cell patch-clamp recordings were performed in either current-clamp or voltage-clamp mode using an EPC-10/2 amplifier controlled by PatchMaster software (HEKA, Germany). The extracellular solution contained (in mM): NaCl 150, KCl 2, CaCl_2_ 2, MgCl_2_ 1, HEPES 10, glucose 10, and glycine 0.01, adjusted to pH 7.3 with NaOH and an osmolarity of 320 mOsm. The intracellular pipette solution contained (in mM): K-gluconate 118, KCl 30, NaCl 5, CaCl_2_ 0.3, EGTA 1, MgATP 2, GTP 0.3, and HEPES 10, adjusted to pH 7.3 with KOH and an osmolarity of 290 mOsm. In some voltage-clamp experiments, potassium salts were replaced with cesium salts, and 3–5 mM QX-314 (an intracellular sodium channel blocker) was added to suppress sodium currents.

Patch electrodes were fabricated from borosilicate glass capillaries and pulled to a resistance of 4– 6 MΩ. Membrane voltage and transmembrane currents were low-pass filtered at 3 kHz and digitized at 10 kHz. Recordings were excluded if leak currents exceeded 200 pA or if series resistance was greater than 30 MΩ.

Protocols for patching and classification of cultured DRG neurons to record T-type calcium currents and to distinguish between neuronal subtypes were performed in extracellular solution containing (in mM): NaCl 150, KCl 2.5, CaCl_2_ 2, MgCl_2_ 1, HEPES 10, and glucose 10, adjusted to pH 7.3 with NaOH and an osmolarity of 325–330 mOsm. For recordings of Ba^2+^ T-currents, the extracellular solution was replaced with (in mM): tetraethylammonium (TEA)-Cl 165, HEPES 10, and BaCl_2_ 2, adjusted to pH 7.4 with TEA-OH and an osmolarity of 305–315 mOsm.

The intracellular pipette solution contained (in mM): K-methanesulfonate 135, KCl 10, EGTA 0.2, MgATP 4, Na-GTP 0.4, Na_2_-phosphocreatine 5, and HEPES 10, adjusted to pH 7.3 with KOH and an osmolarity of 300–305 mOsm. Membrane voltage and transmembrane currents were low-pass filtered at 3 kHz and digitized at 20 kHz. Recordings were excluded if leak currents exceeded 200 pA or if series resistance was greater than 30 MΩ. All experiments were conducted at room temperature.

Liquid junction potentials between the internal KCl/K-methanesulfonate solution and the external NaCl-based solution were calculated as 9.7 mV, and as 15.4 mV between the same internal solution and the TEA-Cl-based external solution. Junction potentials were calculated using the LJCalc program (https://swharden.com/software/LJPcalc/) and were not compensated during recordings.

For activation of total “signature” Na^+^ and K^+^ voltage-gated currents, to classify DRG neurons according their subtypes, depolarizing steps were applied from a holding potential of −100 mV (3.5 s) to test potentials ranging from −60 to +40 mV (200 ms duration) in 20 mV increments. For activation of T-type currents, depolarizing steps were applied from a holding potential of −100 mV (3.5 s) to test potentials ranging from −80 to 0 mV (250 ms duration) in 10 mV increments.

### Estimating the effect of DsiRNA on the functional expression of the Cav3.2 T-channels via recording of intracellular Ca^2+^ concentration

Recordings of free cytoplasmic calcium ion concentration in hippocampal neurons were used to assess the effect of DsiRNA on the functional expression of Cav3.2 T-currents. Neurons in primary culture were co-transfected with DsiRNA and a plasmid encoding CFA. On the second to third day after transfection, calcium transients were measured by loading cells with the fluorescent dye Fluo-5F and applying membrane stimulation protocols in either current-clamp or voltage-clamp mode under whole-cell configuration.

Neurons were stimulated using a patch-clamp micropipette, both under control conditions and in the presence of 10 μM TTA-P2, a selective T-type calcium channel blocker. Whole-cell recordings were performed with micropipettes of 2.5–3 MΩ resistance, which were also used to introduce Fluo-5F dye (200 μM) into the cytoplasm. During experiments, 10 μM nifedipine was included in the external solution to block L-type calcium currents.

Stimulation consisted of a train of 25 action potentials delivered from a holding potential of −90 mV in current-clamp mode at 50 Hz. Each action potential was evoked by a 1 nA, 5 ms current pulse. Calcium transients were recorded as increases in Fluo-5F fluorescence in dendrites, with recordings acquired at 2 Hz.

To minimize errors associated with time-dependent changes in calcium transients (e.g., alterations in intracellular buffering capacity during patch clamp), the amplitude of calcium transients evoked by action potential trains was normalized to the amplitude of transients evoked by a prolonged depolarization (250 ms to +30 mV from a holding potential of −60 mV, voltage-clamp mode). Because of the inactivation properties of T-type channels, their contribution to calcium influx during such prolonged depolarization is minimal. Thus, in cells with significant T-current, the normalized ratio decreases in the presence of TTA-P2, whereas in cells lacking T-current the ratio remains unchanged, even if absolute calcium transient amplitudes vary during the experiment.

## Results

To ensure expression of siRNA capable of suppressing mRNA encoding the Cav3.2 isoform of T-type calcium channels, we have developed and validated several genetic constricts. The constructs were designed to produce highly effective siRNAs in vivo through the cellular processing of short hairpin RNAs (shRNAs) by the endoribonuclease Dicer (Siolas et al., 2005; Maniataki and Mourelatos, 2005). For efficient recognition by Dicer, shRNAs must contain a stem of 25–30 base pairs and two nucleotides protruding at the 3′ end beyond the complementary pairing site (Siolas et al., 2005) (Fig. 2a). Furthermore, the ribonucleotide sequence of the shRNA stem corresponding to the guide strand of the siRNA must be fully complementary to the target site on the Cav3.2 mRNA to enable effective binding and subsequent degradation (Hutvágner and Zamore, 2002) (Fig. 1). Even with complete complementarity, suppression of mRNA expression may be hindered by the formation of internal secondary structures, such as hairpins within the guide strand of shRNA or siRNA. Therefore, it is necessary to experimentally evaluate multiple shRNA constructs targeting different sites to identify the most effective sequence. Direct testing of shRNAs requires expression from viral vectors, which is time-consuming and resource-intensive. As a practical alternative, we initially employed Dicer-substrate siRNAs (DsiRNAs) designed to recognize the same target sites. These molecules are processed by Dicer into functional siRNAs in vivo and can be synthesized and tested more rapidly and cost-effectively in vitro. Such DsiRNAs can be readily synthesized and directly transfected into primary hippocampal or DRG neuron cultures to evaluate their effectiveness in suppressing Cav3.2 mRNA expression.

**Figure 1.**
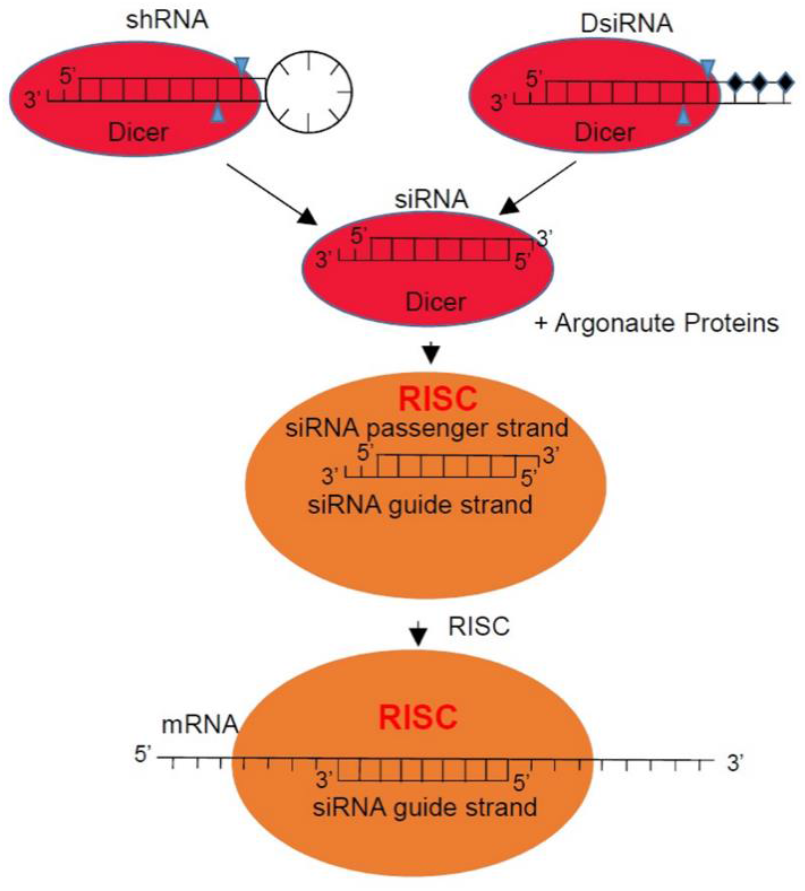
Involvement of shRNA (short hairpin RNA) and DsiRNA (Dicer-substrate siRNA) in mRNA (messenger RNA) degradation. The Dicer endonuclease effectively binds and modifies shRNA and DsiRNA if they have single-stranded overhangs of two ribonucleotides at their 3’ end. To protect against endogenous RNases, the opposite “blunt” 3’ end of the complementary oligonucleotide in the DsiRNA duplex must be protected by modification – replacement of the last three ribonucleotides from the 3’ end with deoxyribonucleotides. (shown by diamonds). With a stem length of shRNA 25-30 nucleotide pairs (n.p.), or a length of the duplex part of DsiRNA 25-27 n.p. Dicer is able to effectively modify these RNAs by cutting the duplex part of the RNA at a distance of 21 b.p. from the single- stranded 3’ end of these RNAs (the sites of cuts are shown by blue triangles) with the formation of shorter siRNAs, which it loads into the blocking complex RISC (RNAi-induced silencing complex) the main components of which are Argonaute proteins. Single-stranded overhangs at the 3’ ends of shRNA and DsiRNA play a decisive role in their binding to Dicer and loading of the newly formed siRNA into RISC. After loading into RISC, the complementary strand of siRNA (complementary to the strand that formed the protruding 3’ end of the duplexes) is cleaved by Ago2 protein, and the guide strand of siRNA (the one that formed the protruding 3’ end of the duplexes) binds to the complementary region of mRNA (target) and, thus, activates the degradation of this mRNA by RISC proteins.

**Figure 2.**
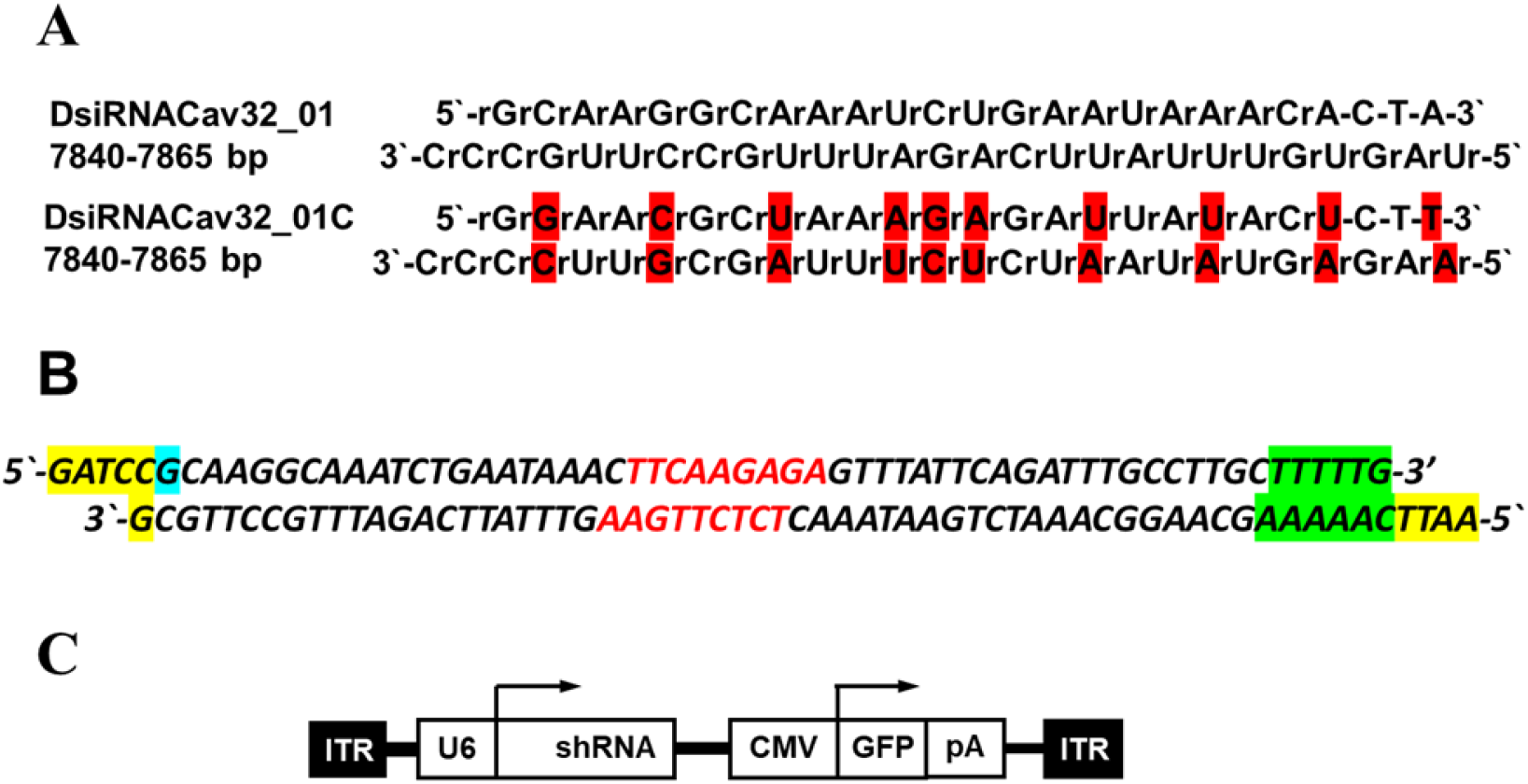
Structure of **(A)** DsiRNA, **(B)** duplex for cloning the shRNA coding region between BamHI and EcoRI sites under the U6 promoter as part of the expression cassette, **(C)** expression cassette with inverted terminal repeats ITRs required for packaging into AAV6 viral envelopes.

### Designing DsiRNA and shRNA-coding constructs for interference with the mRNA of the Cav3.2 isoform of the T-type channels

Designing DsiRNAs and shRNAs was necessary for supressing translation of the Cav3.2 alpha subunit of the T-type channels in nociceptive DRG neurons in order to downregulate pain symptoms of the PDN. Both shRNAs and DsiRNAs are effectively processed by endoribonuclease Dicer if they have 25-30 bp long stem or duplex part and two bp long overhangs at their 3’-ends (Fig. 1). We have also taken additional measures for fidelity of the cut of shRNA by endoribonuclease Dicer and preferred loading of the guiding strand into RISC – all for the sake of specificity of the target mRNA knockdown and minimization of the nonspecific activity of RISC induced by shRNA on the other genes (Bofill-De Ros and Gu, 2016). Such measures included: the length of a stem duplex was taken equal to 21 b.p., the length of the loop was taken longer than 6 b.p., two bp long overhangs at their 3’- ends (Fig. 1) were introduced for fidelity of the Dicer cut. The first ribonucleotide on the 5’-end of the guiding strand after Dicer-made cut were chosen to be rA ot rT for effective binding of Ago2-protein and preferred loading of this strand into RISC. Additionally, target sequence for DsiRNA and shRNA was chosen in the 3’- untranslated AT-rich region for easy unwinding of mRNA after seed-sequence is annealed. First, we have designed DsiRNA with its target sequence in 3’-untranslated region of the Cav3.2 mRNA (Fig. 2A) to check its effectiveness in interfering with the expression of the T-channels in the neurons of primary cell culture (hippocampal cultures or DRG neurons culture). After that, we used its target sequence to design shRNA stem-coding sequence (Fig. 2B). The whole coding sequence for the shRNA was cloned in the vector pAAV6-sh[control] purchased from the Addgene (USA), in the body of the expression cassette, under the U6 promotor, flanked with AAV inverted terminal repeats (ITRs) (Fig. 2C) for effective packing into AAV particles.

### Selecting and testing DsiRNA for suppression of Cav3.2 T-channel expression in rat hippocampal neurons

To identify potential DsiRNAs capable of blocking expression of rat Cav3.2 T-type calcium channels, we used the Dicer-substrate siRNA design tool provided by Integrated DNA Technologies (IDT, Coralville, Iowa, USA; https://eu.idtdna.com/site/order/tool/index/DSIRNA_CUSTOM). Target sites were selected from the nucleotide sequence of the Cav3.2 α-subunit mRNA (GenBank reference NM_153814.2, Cacna1h, Rattus norvegicus). Among the regions identified by the program, three sites were chosen: one located in the 3′-untranslated region (positions 7840–7865 bp), and two located within the coding region (positions 6955–6980 bp and 4312–4337 bp). Duplex DsiRNAs designed against these sites were predicted to be stable in transfected cells and to effectively suppress Cav3.2 expression (Fig. 2A).

Based on the selected targets the program has designed duplex DsiRNAs that should be stable in transfected cells and effectively suppress the expression of Cav3.2 T-channels (Fig. 2A). Based on the selected sequences, DsiRNAs were ordered from IDT and dissolved on arrival in water at concentration sutable for contransfection with CFP-expressing plasmid into neurons of the primary hippocampal or DRG neurons cell culture. As mentioned in a section Materials and Methods devoted to designing of DsiRNA this step was used to find effective target sequence in the mRNA primary structure to include it in a structure of the shRNA on a second step, to be expressed in a plasmid vector for further assembly into AAV viral particles. At the same time DsiRNA have its own value as a therapeutic medication (drug) as it is designed in a way to protect it from RNA nucleases (prolong its half-decay time *in vivo*) and can be delivered in *vivo* to primary nociceptive DRG neurons using certain transfection reagents.

DsiRNA variant DsiRNACav32_01 (Fig. 2A) was developed to target 3’-untranslated region of the Cav3.2 T-type channels mRNA to interfere with its expression. It was cotransfected with the plasmid expressing cyan fluorescent protein (CFP) into the neurons of hippocampal cell culture and was proved to be effective in inhibiting the Ca^2+^ concentration transients (calcium transients) inside neuron’s dendrites mediated by T-type channels during trains of APs induced by corresponding current-clamp stimulations (Fig. 3). In transfected neurons, no statistically significant change in the ratio of calcium transient amplitudes, calculated as described in Materials and Methods, was detected upon application of TTA-P2, with an increase of 2.7% +/-1.9% (p>0.05, 9 cells), that should indicate the absence of T-current in such neurons. At the same time, in control untransfected cells (control cells were selected for size and morphology similar to transfected cells, on the same slide as the original cells) after application of TTA-P2, this ratio decreased by 13% +/- 4% (p < 0.005, 9 cells). We also performed similar experiments with a control DsiRNA duplex, DsiRNACav32_01C, that was as similar as possible to the DsiRNACav32_01, but had mismatch mutations to not block T-channel expression (Fig. 2A). With the DsiRNACav32_01C duplex in transfected neurons, this ratio decreased by 13% +/- 3% (p < 0.005, 14 cells; Fig 4A). For control untransfected cells in this series of experiments, the ratio decreased by 12%+/- 4% (p < 0.005, 14 cells), when TTA-P2 was used (Fig. 4B).

**Figure 3.**
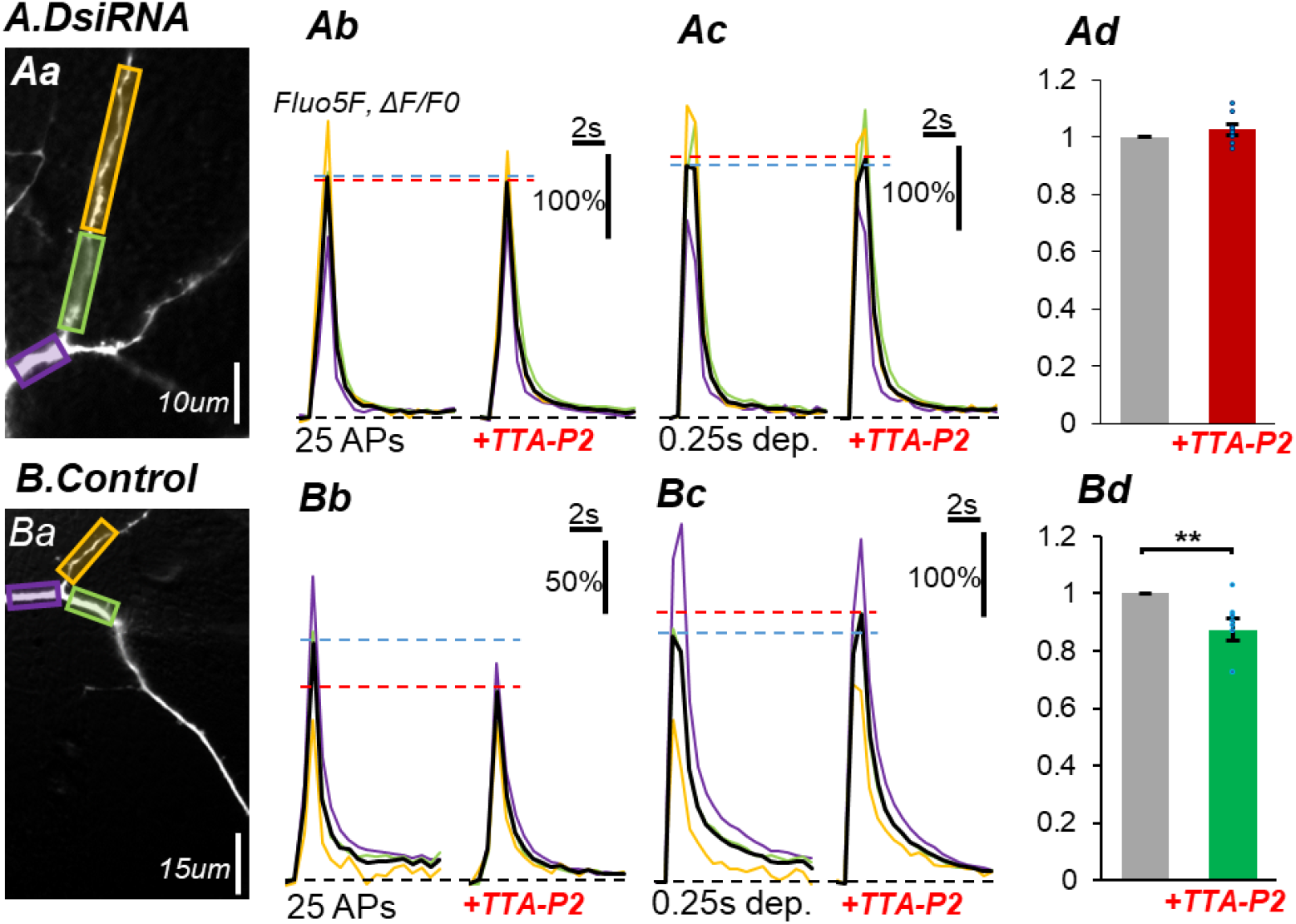
Comparison of calcium transients in primary hippocampal neurons cotransfected with DsiRNACav32_01 and a plasmid expressing the fluorescent protein CFP **(A)** and in naive neurons transfected with a plasmid expressing the CFP protein alone **(B). Aa, Ba** – Fluo5F fluorescent image of the dendritic tree of neurons transfected with DsiRNACav32_01 **(Aa)** and naive neurons **(Ba). Ab, Bb, Ac, Bc** – Fluo5F calcium dye fluorescence spike amplitude upon stimulation of neurons with sequences of 25 action potentials in neurons transfected with DsiRNACav32_01 and a plasmid encoding CFP **(Ab left)** and control neurons transfected with only a plasmid encoding CFP **(Bb left)**, followed by addition of a specific Cav3.2 T-channel blocker - TTA-P2 to the experimental chamber **(Ab right** - for DsiRNACav32_01 and **Bb right** - for control), or upon stimulation of neurons by membrane depolarization from -90 mV to +30 mV for 250 msec: **(Ac left)** - DsiRNACav32_01 without TTA-P2, **(Bc left)** - naive neurons without TTA-P2, **(Ac right)** - DsiRNACav32_01 + TTA-P2, **(Bc right)** - naive neurons + TTA-P2. **Ad, Bd** – Comparison of the amplitude of the fluorescence spike of the calcium dye Fluo5F, calculated by normalizing the amplitude measured on **(Ab)** to that measured on **(Ac)** or that measured on **(Bb)** to that measured on **(Bc)**, correspondently, before application of TTA-P2 **(Ad** or **Bd, column on the left**) and after application **(Ad** or **Bd, column on the right)**.

**Figure 4.**
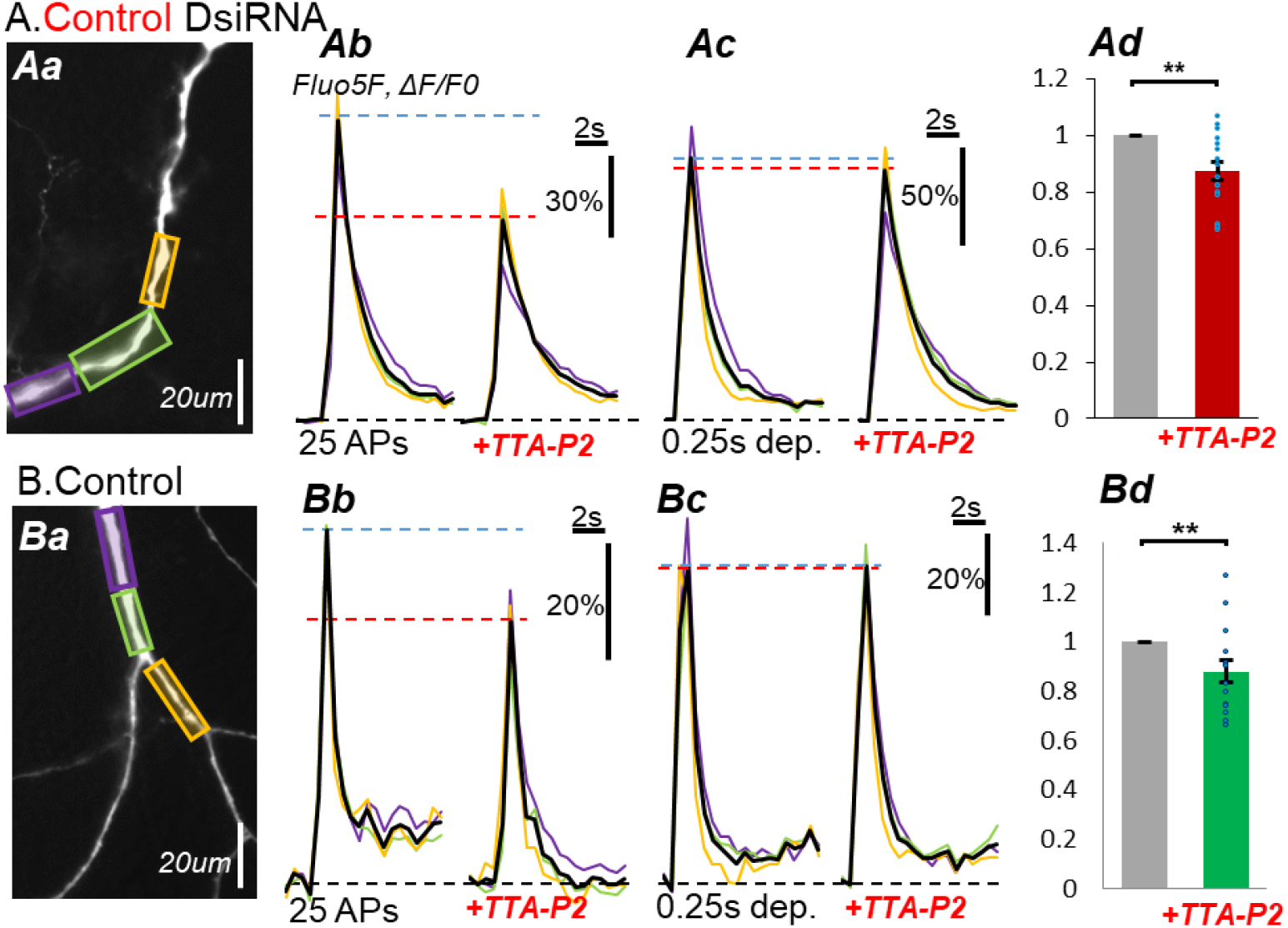
Comparison of calcium transients in primary hippocampal neurons cotransfected with the control mutated DsiRNA variant, DsiRNACav32_01C, and a plasmid expressing the fluorescent protein CFP **(A)** and in naive neurons transfected with a plasmid expressing the CFP protein alone **(B). Aa, Ba** – Fluo5F fluorescent image of the dendritic tree of neurons transfected with DsiRNACav32_01C **(Aa)** and naive neurons **(Ba). Ab, Bb, Ac, Bc** – Fluo5F calcium dye fluorescence spike amplitude upon stimulation of neurons with sequences of 25 action potentials in neurons transfected with DsiRNACav32_01C and a plasmid encoding CFP **(Ab left)** and control neurons transfected with only a plasmid encoding CFP **(Bb left)**, followed by addition of a specific Cav3.2 T- channel blocker - TTA-P2 to the experimental chamber (**Ab right** - for DsiRNACav32_01C and **Bb right** - for naïve neurons), or upon stimulation of neurons by membrane depolarization from -90 mV to +30 mV for 250 msec: **(Ac left)** - DsiRNACav32_01C without TTA-P2, **(Bc left)** - naive neurons without TTA-P2, **(Ac right)** - DsiRNACav32_01C + TTA-P2, **(Bc right)** - naive neurons + TTA-P2. **Ad, Bd** – Comparison of the amplitude of the fluorescence spike of the calcium dye Fluo5F, calculated by normalizing the amplitude measured on **(Ab)** to that measured on **(Ac)** or that measured on **(Bb)** to that measured on **(Bc)**, correspondently, before application of TTA-P2 **(Ad** or **Bd, column on the left)** and after application **(Ad** or **Bd, column on the right)**.

Taken together, these results demonstrate that DsiRNACav32_01 effectively and specifically suppresses Cav3.2 T-currents in cultured hippocampal neurons. The near-complete blockade of T-type channel activity by this construct highlights its potential as a candidate for therapeutic development in the treatment of painful diabetic neuropathy.

### Designing and Testing an shRNA-Expressing Construct Targeting Cav3.2 T-Type Channels in DRG Neurons

To generate a long-lasting RNA interference tool, the target sequence identified as effective in DsiRNA experiments was incorporated into the stem structure of an shRNA. Specifically, the sequence was used as the “passenger” strand, complementary to the “guide” strand designed to bind the Cav3.2 mRNA target site. Compared to DsiRNAs, shRNAs offer the advantage of sustained interference when expressed from a promoter within a suitable vector, and they can be selectively delivered to specific neuronal populations when packaged into adeno-associated virus (AAV) particles. For this purpose, we employed the pAAV-sh[control] vector (Addgene, USA; map available at https://www.addgene.org/75438/sequences/), which contains an expression cassette flanked by AAV inverted terminal repeats (ITRs). A duplex DNA fragment encoding the shRNA was cloned into the vector using BamHI and EcoRI restriction sites (Fig. 2B, C).

The resulting plasmid, pAAV-shRNA::Cav3.2-N1DF, was tested for its ability to suppress T-type currents in primary DRG neuron cultures. Transfected cells were identified by green fluorescent protein (GFP) expression encoded by the vector, while non-fluorescent cells served as untransfected controls. A mutated shRNA sequence (NegD) was used as a negative control. Whole-cell voltage-clamp recordings were performed using extracellular solutions containing Ba^2+^ and TEA-Cl. The presence of low-voltage activated Ba^2+^ currents, with activation thresholds between −60 and −40 mV, was taken as evidence of T-type channel activity. Specificity for Cav3.2 channels was confirmed by application of 50 μM Ni^2+^, which selectively inhibits this isoform.

All neurons transfected with pAAV-shRNA::Cav3.2-N1DF exhibited Ba^2+^ current activation thresholds above −40 mV (typically between −30 and −20 mV), consistent with the absence of low-voltage activated T-type currents (Fig. 5, Ad, Bd, Cd, left). In contrast, the majority of untransfected neurons displayed low-voltage activated currents that were inhibited by Ni^2+^ (Fig. 5, Ad, Bd, Cd, middle), confirming Cav3.2 channel involvement. The mean activation threshold of Ba^2+^ currents in transfected neurons was significantly higher than in untransfected controls (−28.9 ± 3.0 mV, n = 9 vs. −52.2 ± 2.8 mV, n = 9; p < 0.001), indicating robust suppression of T-type currents (Fig. 5, D column N1DF vs. Not trans.).

**Figure 5.**
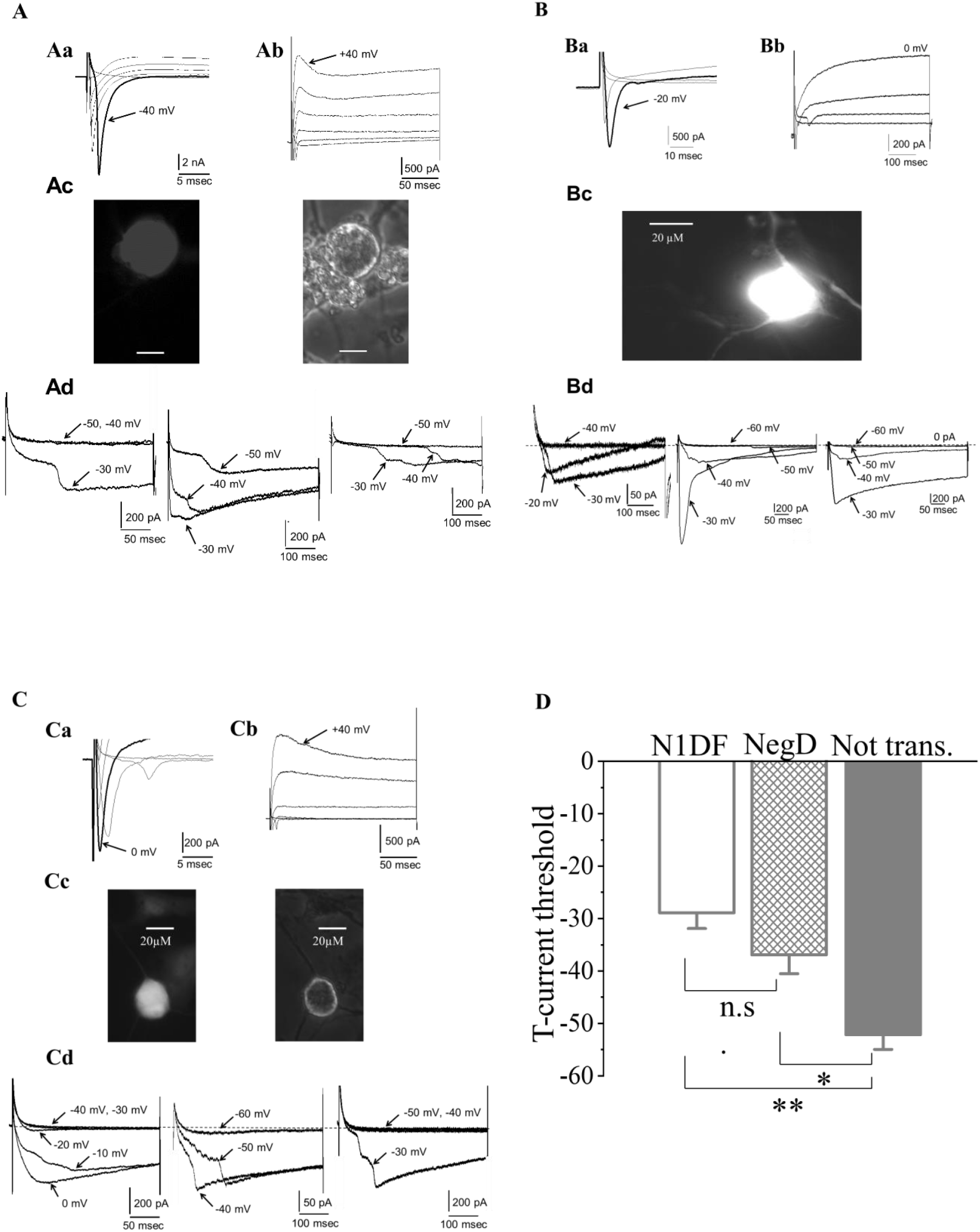
shRNA variant N1DF developed for knockdown of T-type Ca^2+^ channels produced significant suppression of channel functioning. **A, B and C** demonstrate a suppression of T-current in three different types of DRG neurons. Na^+^-currents **(Aa, Ba, Ca)**, K^+^-currents **(Ab, Bb, Cb)** and cell phenotype characteristics **(Ac, Bc, Cc). Ac, Bc, Cc** - images of representative DRG neurons transfected with the plasmid pAAV-shRNA::Cav3.2-N1D containing N1DF variant of shRNA. **(Ac, Bc, Cc left -** GFP fluorescent images and **Ac, Bc, Cc right -** phase contrast images**). Ad, Bd, Cd -** comparison of Ba^2+^ currents in the transfected neurons **(left)**, control untransfected neurons of the same subtype **(center)** and the neurons transfected with the negative control (NegD) plasmid variant pAAV-shRNA::Cav3.2-NegD **(right)**. Patterns of Na^+^-current and K^+^-current were recorded at depolarization from -100 mV to -60 mV through +40 mV with a step 20 mV. Presented Ba^2+^ currents **(Figures (Ad, Bd, Cd) left, center and right)** were recorded at depolarization steps from -100 mV to the potentials indicated by the respective arrows. **(D)** demonstrates the pooled statistics in respect of Ba^2+^ current thresholds (9 transfected cells with the inhibiting plasmid (N1DF) from 9 rats, 9 untransfected cells from 9 rats and 7 transfected cells with the negative (noninhibiting) control plasmid (NegD) from 7 rats). The mean threshold for the N1DF-transfected neurons (-28.9 + 3.0 mV) was significantly higher than one for the untransfected neurons (-52.2 + 2.8 mV, p<0.001) and was insignificantly higher than for the negative control NegD-transfected neurons (-37.1 + 3.6 mV, p>0.1)

The negative control construct NegD produced mixed results. Of seven neurons transfected with NegD, three retained low-threshold T-type currents, while four did not. However, three of the four neurons lacking T-currents belonged to DRG subtypes not previously characterized in our studies as ones having T-type current and are likely to lack Cav3.2 expression. These neurons were excluded from statistical analysis. Comparison of the proportion of neurons losing T-currents after transfection with N1DF versus NegD revealed a significant difference between groups (p = 0.014, Fisher’s exact test).

Together, these findings demonstrate that the shRNA-expressing construct pAAV-shRNA::Cav3.2-N1DF effectively interferes with Cav3.2 mRNA expression, leading to suppression of low-threshold T-type currents in nociceptive DRG neurons.

## Discussion

As highlighted in the Introduction, numerous studies have causally linked the functional overexpression of Cav3.2 T-type calcium channels in nociceptive dorsal root ganglion (DRG) neurons with painful diabetic neuropathy (PDN). In vivo experiments using antisense oligonucleotides or selective pharmacological blockers have confirmed that suppression of Cav3.2 channel expression or activity alleviates painful symptoms in PDN models (Latham et al., 2009; Messinger et al., 2009). Despite these findings, practical approaches for long-term, cell-specific inhibition of Cav3.2 channels suitable for clinical application have not yet been developed.

In this study, we designed and validated a plasmid construct based on an adeno-associated viral (AAV) vector that enables transcription of shRNA targeting Cav3.2 channels from a constitutive promoter. This expression cassette can be packaged into AAV6 particles, which display tropism for nociceptive DRG neurons, thereby providing a strategy for long-lasting inhibition of Cav3.2 channels in vivo. The design of the shRNA construct incorporated current knowledge of structural features required for efficient Dicer recognition and processing, including stem length, loop size, and the presence of 3′ overhangs, as well as preferential loading of the guide strand into the RNA-induced silencing complex (RISC) via Ago2 (Bofill-De Ros and Gu, 2016). To validate target sequence selection, DsiRNAs with identical sequences were designed and tested in vitro, ensuring that the chosen shRNA would be effective when expressed from the plasmid.

Data obtained from hippocampal primary cultures confirmed the specificity and reliability of the DsiRNA constructs. Neurons transfected with functional DsiRNA were compared both to untransfected controls and to neurons transfected with mutated, nonfunctional DsiRNA. The use of the Cav3.2-specific blocker TTA-P2 further demonstrated specificity: neurons transfected with functional DsiRNA showed no additional effect of TTA-P2 compared to untransfected controls, confirming that the DsiRNA acted specifically on Cav3.2 channels.

In DRG primary cultures, the plasmid construct pAAV-shRNA::Cav3.2-N1DF effectively suppressed functional expression of low-threshold T-type currents. Specificity of the N1DF shRNA was confirmed by applying 50 μM Ni^2+^, which selectively inhibits Cav3.2 channels, to untransfected control neurons of the same subtype. In transfected neurons, the threshold for Ba^2+^ current activation shifted significantly toward high-threshold levels compared to untransfected DRG neurons (Fig. 4), indicating reduced T-current expression.

The pathological overexpression of Cav3.2 channels in nociceptive DRG neurons under diabetic conditions is widely recognized as a key mechanism underlying hyperalgesia, allodynia, and spontaneous pain. Our constructs—both DsiRNAs and the pAAV-shRNA::Cav3.2 plasmid— demonstrated effective interference with Cav3.2 expression in vitro in rat hippocampal and DRG cultures. These findings provide a strong rationale for further in vivo testing. DsiRNAs can be injected into the sciatic nerve to directly assess their ability to reduce diabetic neuropathic pain in behavioral assays. However, because synthetic siRNAs have a limited lifespan in vivo (approximately 5–6 days), the shRNA-expressing plasmid offers a more promising long-term solution. If DsiRNAs prove effective in vivo, the corresponding shRNA-expressing vector should be tested for sustained therapeutic effects by direct injection into DRG or sciatic nerve. Ultimately, the expression cassette can be packaged into AAV6 particles for cell-specific delivery to nociceptive DRG neurons, enabling prolonged suppression of Cav3.2 channel expression and potentially providing durable relief of painful symptoms in PDN.

While our findings provide compelling in vitro evidence for the effectiveness of DsiRNA and shRNA constructs in suppressing Cav3.2 expression, several limitations must be acknowledged. First, all experiments were conducted in rodent primary cultures, and translation to in vivo models will be essential to confirm therapeutic efficacy and safety. Second, although AAV6 vectors display tropism for nociceptive DRG neurons, off-target transduction in other cell types cannot be excluded and requires systematic evaluation. Third, long-term expression of shRNA may carry risks of unintended gene silencing or immune activation, which must be carefully assessed in preclinical studies. Finally, while DsiRNAs offer a rapid screening tool and potential therapeutic application, their limited stability in vivo highlights the need for optimized delivery systems or chemical modifications to extend their half-life. Future work should therefore focus on in vivo validation of both DsiRNA and AAV6-shRNA constructs, assessment of off-target effects, and exploration of delivery strategies that maximize specificity and durability.

## Author Contributions

Conceptualization, D.D., O.H. N.V. and P.B.; methodology, D.D., O.H. N.H. A.D., N. K. and P.B.; electrophysiology, digital imaging and culturing, D.D., A.D., N.K. and E.L.; molecular genetics, O.H., N.H., A.R, N.S. and D.D.; validation, O.H., D.D. N.V., N.K. and P.B.; writing, D.D., A.D. and P.B.; supervision, P.B. All authors have read and agreed to the published version of the manuscript.

## Institutional Review Board Statement

The study was conducted according to the guidelines of the Declaration of Helsinki and approved by the Institutional Bioethics Committee of the Bogomoletz Institute of Physiology at NASU.

## Informed Consent Statement

Not applicable.

## Data Availability Statement

The data are available upon reasonable request.

